# GSKB: A gene set database for pathway analysis in mouse

**DOI:** 10.1101/082511

**Authors:** Liming Lai, Jason Hennessey, Valerie Bares, Eun Woo Son, Yuguang Ban, Wei Wang, Jianli Qi, Gaixin Jiang, Arthur Liberzon, Steven Xijin Ge

## Abstract

Interpretation of high-throughput genomics data based on biological pathways constitutes a constant challenge, partly because of the lack of supporting pathway database. In this study, we created a functional genomics knowledgebase in mouse, which includes 33,261 pathways and gene sets compiled from 40 sources such as Gene Ontology, KEGG, GeneSetDB, PANTHER, microRNA and transcription factor target genes, *etc*. In addition, we also manually collected and curated 8,747 lists of differentially expressed genes from 2,526 published gene expression studies to enable the detection of similarity to previously reported gene expression signatures. These two types of data constitute a Gene Set Knowledgebase (GSKB), which can be readily used by various pathway analysis software such as gene set enrichment analysis (GSEA). Using our knowledgebase, we were able to detect the correct microRNA (miR-29) pathway that was suppressed using antisense oligonucleotides and confirmed its role in inhibiting fibrogenesis, which might involve upregulation of transcription factor SMAD3. The knowledgebase can be queried as a source of published gene lists for further meta-analysis. Through meta-analysis of 56 published gene lists related to retina cells, we revealed two fundamentally different types of gene expression changes. One is related to stress and inflammatory response blamed for causing blindness in many diseases; the other associated with visual perception by normal retina cells. GSKB is available online at http://ge-lab.org/gs/, and also as a Bioconductor package (gskb, https://bioconductor.org/packages/gskb/). This database enables in-depth interpretation of mouse genomics data both in terms of known pathways and the context of thousands of published expression signatures.

## INTRODUCTION

Pathway analysis is a key step in analyzing high-throughput genomics data. The goal of pathway analysis is to determine if coherent change in gene expression occurs among a set of genes related to a molecular pathway or biological function. Many methods have been developed to achieve this goal (see review in [1]). While gene set enrichment analysis (GSEA) [2], one of the most popular programs, is based on non-parametric Kolmogorov-Smirnov statistic, several parametric algorithms have been developed [3, 4]. The foundation for all of these algorithms is a database of curated pathways and functional categories. The Molecular Signatures Database (MSigDB) [5] is a collection of gene lists initially developed for GSEA [2]. While its main focus is human, MSigDB also includes a small number of gene sets for mouse and rat. In addition to existing pathway databases, MSigDB also includes lists of differentially expressed genes manually collected from published gene expression studies related to genetic and chemical perturbations. Inclusion of these gene sets enables detection of the co-regulation of genes similar to those reported in the literature. The MSigDB has been widely used and greatly facilitate pathway analysis in human genomics studies.

As much of the work has been focused on gene sets in human, there is an urgent need of comprehensive pathway databases for other model organisms. Some researchers have to convert human genes into mouse orthologs in order to use MSigDB [6]. Other previous efforts include Genetrail [7], which constructs its own database by extracting information from several sources such as Gene Ontology [8], KEGG [9] etc. GeneSetDB is a larger collection of gene sets for several species based on 26 public databases [10]. Pathway and gene-set enrichment database(PAGED), which covers 20 species, derives information from many different sources including GeneSetDB and other disease-gene association data [11]. We recently created a pathway database for Arabidopsis [12].

In this study, we sought to develop a pathway knowledgebase for mouse, an important model organism for the study of many human diseases. We compiled gene sets from a large number of existing annotation databases as well as thousands of primary publications, which covers a wide spectrum of genetic, genomic and biological information. These gene sets forms a foundation for in-depth interpretation of mouse genomics data, supporting the use of mouse as a model for understanding human biology and diseases. We also demonstrate the use of this knowledgebase to generate testable hypotheses through several examples.

## METHODS

Since most gene expression studies deposit gene expression data and publication information in public repositories, we first search for publications in Gene Expression Omnibus (GEO, www.ncbi.nlm.nih.gov/geo/) and ArrayExpress (www.ebi.ac.uk/arrayexpress/). We focused on mouse related expression studies in this study. Based on the information, the full text of the papers and their supplementary materials were retrieved. Then gene lists were compiled and curated after reading the papers and their supplementary materials. Similar to MSigDB [5] and AraPath [12], an unique identifier was assigned to each gene list. The names often start with the last names of the first authors and end with “_up”, or “_down” to indicate whether the genes are up-regulated or down-regulated, respectively. A brief one-sentence description was also given to each gene list. For example, gene list VENEZIA_FETAL-LIVER_UP represents genes highly expressed in fetal liver. A long description for the gene lists are abstracts of the paper from PubMed. We retrieved and processed all papers linked to GEO and ArrayExpress that we can found at the time of the study. We tried to collect all lists of differentially expressed genes reported in the literature. Larger lists (>3000 genes) are excluded. Finally, all gene lists were merged into an Excel spreadsheet to be further processed. One key step is the conversion of various gene IDs from different sources to NCBI gene symbols and Mouse Genome Informatics (MGI) IDs. The conversion was made based on the mouse genes information at NCBI, including platform definition files at GEO and other gene information files such as gene2accession.gz, Mm.data.gz, Mm.gb_cid_lid, and All_data.gene_info.gz (ftp://ftp.ncbi.nih.gov). Conversion to MGI IDs is carried out using information from MGI web site (www.informatics.jax.org). A Perl program was created to convert these gene IDs.

Gene lists from existing sources were manually imported in July 2013. Our web site (http://gskb.ge-lab.org) was assembled using a combination of the Python programming language, Django web framework, MySQL, JavaScript, and html. More detailed information about GSKB is included in supplementary file 3 which follows the BioDBcore guidelines [13]. This web site also includes data for other species such as the Arabidopsis gene sets [12], as well as other species that are still in the process.

Gene expression datasets were downloaded from GEO with accession numbers GSE40261and GSE27035. The raw Affymetrix .CEL files were processed using robust multiarray analysis (RMA) algorithm [14] as implemented in Bioconductor [15]. “Present” or “Absent” calls were calculated using the Affymetrix MAS5 algorithm based on Wilcoxon Rank Sum test. Probe-sets with “Absent” calls across all samples were deemed not significantly above background and removed from further analysis. Probe-set IDs were mapped to official gene symbol based on the mapping in Bioconductor package mouse4302.db. If multiple probe-sets were mapped to the same gene, we retained the one with the largest standard deviation. The processed gene expression data is available as supplementary file 4. GSEA version 2.0 was used with default parameters to analyze GSE40261. We also used another method, parametric analysis of gene set enrichment (PAGE) [3], implemented in the PGSEA package [16]. Gene expression data was mean-centered before using PGSEA, which calculates a *t*-statistic for each gene-set in every sample to measure whether the genes are collectively up- or down-regulated. The *t*-statistics were used in an analysis of variance (ANOVA) to test whether there is significant difference between sample groups. Gene sets are ranked by the difference in average *t*-statistic.

## RESULTS AND DISCUSSION

GSKB is a comprehensive knowledgebase for pathway analysis in mouse. It complements MSigDB, which contains a small number of mouse gene lists. Unlike GeneSetDB [10], GSKB includes a large number of lists of differentially expressed genes. The data can be downloaded and used by various software packages for pathway analysis. Our web site also includes an interface for keyword search. The keywords can be a topic-related (“stem cell”), or a gene symbol (“SOX2”), which will be compared against all the descriptions including the abstracts as well as gene lists. In addition, we also provide a similarity search page, where users can upload a gene list, and our web server will compare user’s gene lists with all the gene lists in the database and return those that have statistically significant overlaps.

### Construction of the knowledgebase

As a first step, we gathered annotation information from 40 existing databases for mouse-related gene sets (Table 1). These gene sets are divided into 8 categories, namely, Co-expression, Gene Ontology, Curated pathways, Metabolic Pathways, Transcription Factor (TF) and microRNA target genes, location (cytogenetics band), and others. We used information in GeneSetDB [10] for some of the databases. Detailed information on these 40 sources and the citations is available in supplementary file 1.

**Table 1.**
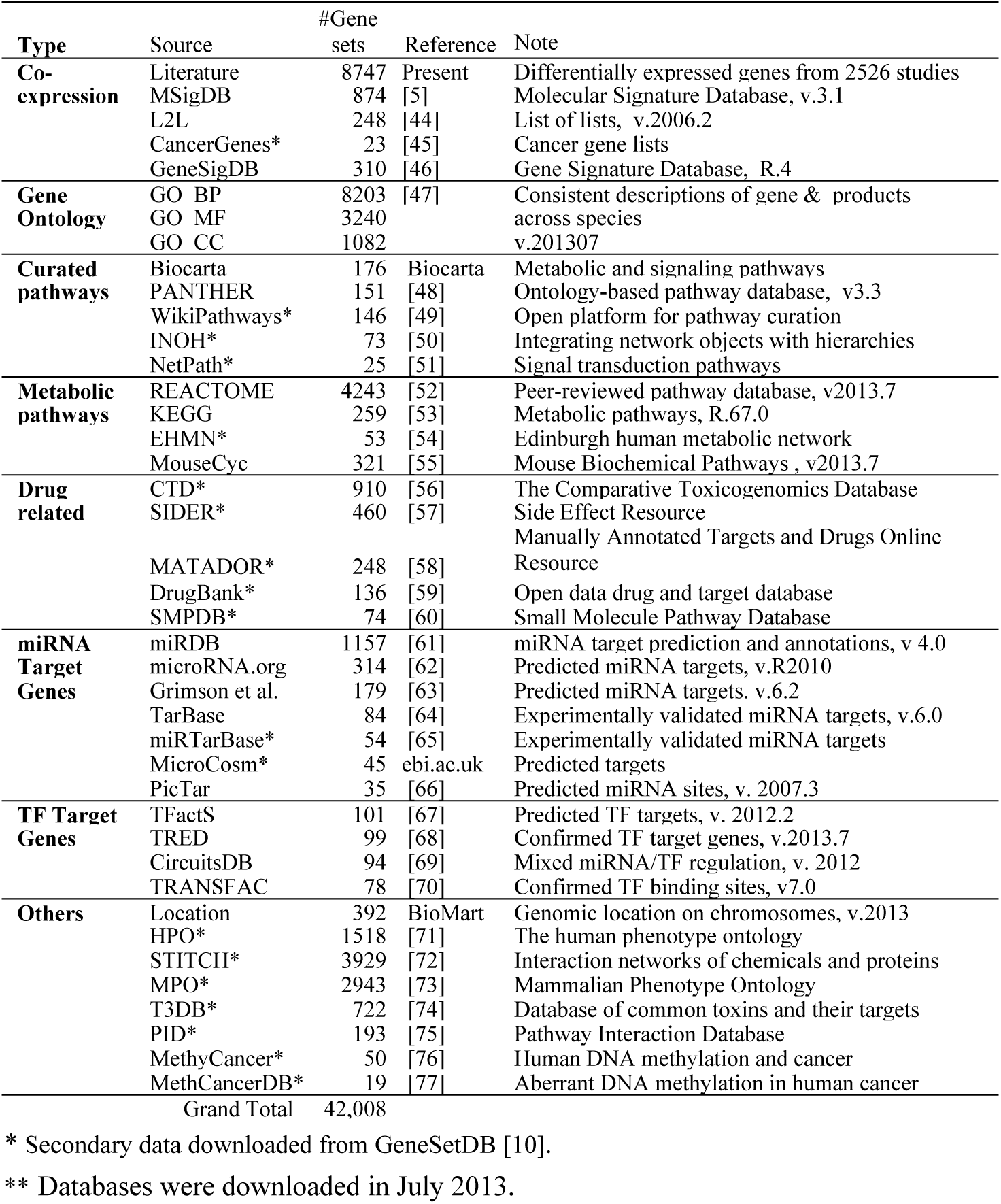
Sources for gene sets in GSKB.

The gene lists from literature were retrieved manually from individual gene expression studies through a process similar to the one used to create AraPath, a similar resource for Arabidopsis[12]. As most expression studies upload raw data to repositories like GEO and ArrayExpress, we used the meta-data in these databases to search for publications. We scanned all datasets we can found and retrieved 4,313 potentially useful papers reporting gene expression studies in mouse. These papers were individually read by curators to identify lists of differentially expressed genes in various conditions. We compiled a total of 8,747 lists of differently expressed genes from 2,518 of papers (See supplementary information 2 for citations). Each gene list was annotated with a unique name, brief description, and publication information, similar to the protocol used in MSigDB and Arapath [12]. These gene lists constitute a large collection of published expression signatures that form a foundation for interpret new gene lists and expression profiles.

These 8,747 gene lists collected from literature include a total of 29,876 unique gene symbols. Interestingly, the distribution of the sizes of published gene lists approximates a normal distribution on logarithm scale (Fig. 1). Although the median size is 51, there are many gene lists containing a few hundreds of genes. The most frequently appeared genes in these lists are shown in Table 2. It includes genes related to cell cycle (CCND2, CDKN1A), and immune response (SOCS3, FOS). Many were intensively-studied, as suggested by the number of related PubMed hits when using gene symbol in keyword searches.

**Fig. 1.**
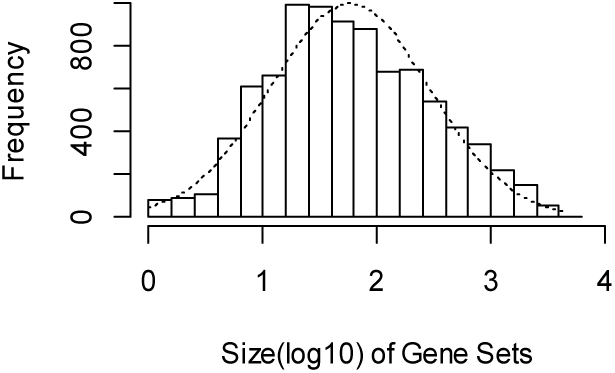
Size distribution of the number of differentially expressed genes reported in the literature

**Table 2.** Top 10 most frequent genes in 8,747 published lists of differentially expressed genes.

| Freq-<br>uency | Gene<br>Symbol | Official Full Name | #PubMed<br>Citations |
| --- | --- | --- | --- |
| 615 | CDKN1A | cyclin-dependent kinase inhibitor 1A (P21) | 660 |
| 498 | CCND2 | cyclin D2 | 239 |
| 496 | MT1 | metallothionein 1 | 272 |
| 493 | SOCS3 | suppressor of cytokine signaling 3 | 262 |
| 490 | ID2 | inhibitor of DNA binding 2 | 256 |
| 477 | EGR1 | early growth response 1 | 332 |
| 465 | IGF1 | insulin-like growth factor 1 | 668 |
| 463 | GJA1 | gap junction protein, alpha 1provided | 602 |
| 457 | SCD1 | stearoyl-Coenzyme A desaturase 1 | 156 |
| 454 | FOS | FBJ osteosarcoma oncogene | 440 |

### Using the database

The database can be downloaded at our web site (http://ge-lab.org/gs/), and used with various pathway analysis software. The web site can also be searched using keywords representing either gene names or pathway names. Using the “Find related genesets” page, users can also upload their own lists of genes and search the database for gene sets with significant overlap. Finally, we also created a Bioconductor package “gskb” (https://bioconductor.org/packages/gskb/) that provide access to this data.

### In-depth analysis of the expression profile of miR-29 silencing and induction

To test if known pathways could be detected, we re-analyzed a gene expression dataset from Hand et al. [17], which is available in GEO database with accession number GSE40261. In this experiment, mice were injected of antisense oligonucleotides against miR-29a or a scrambled sequence as control. The antisense oligonucleotides should suppress the expression of miR-29a, thereby affect the expression of the set of genes inhibited by this microRNA in liver tissue. We downloaded and processed the raw Affymetrix files and used different pathway analysis software to detect significantly altered pathways (See Methods for details). We first focused on 1869 predicted microRNA target gene sets. Table 3 shows the top gene sets using GSEA. It is clear that the most significantly affected pathways are those related to miR-29a, b, and c. We also used another method, parametric analysis of gene set enrichment (PAGE) [3], as implemented in the PGSEA package [16]. The results are shown in Fig. 2. The top 6 pathways are all miR-29 target gene sets. Therefore, the expected molecular pathway was detected using different software.

**Table 3.** Significant gene sets from pathway analysis using GSEA. ES: Enrichment Score, NES: Normalized Enrichment Score, P-val: Nominal P values, FDR: False discovery rate, FWER: family-wise error rate.

| NAME | SIZE | ES | NES | P-val | FDR | FWER |
| --- | --- | --- | --- | --- | --- | --- |
| MIRNA_MM_WANG_MMU-**MIR-29C**-3P | 220 | 0.604 | 1.722 | 0 | 0.013 | 0 |
| MIRNA_MM_WANG_MMU-**MIR-29A**-3P | 217 | 0.608 | 1.715 | 0 | 0.013 | 0.013 |
| MIRNA_MM_WANG_MMU-**MIR-29B**-3P | 221 | 0.591 | 1.711 | 0 | 0.013 | 0.013 |
| MIRNA_MM_WANG_MMU-MIR-767 | 112 | 0.558 | 1.559 | 0 | 0.099 | 0.262 |
| MIRNA_MM_WANG_MMU-MIR-5620-5P | 25 | 0.657 | 1.528 | 0.028 | 0.171 | 0.392 |
| MIRNA_MM_WANG_MMU-MIR-3065-3P | 192 | 0.431 | 1.519 | 0 | 0.163 | 0.428 |
| MIRTARBASE_MM_HSA-**MIR-29B** | 27 | 0.805 | 1.507 | 0.025 | 0.162 | 0.458 |
| MIRTARBASE_MM_HSA-**MIR-29A** | 21 | 0.763 | 1.503 | 0 | 0.158 | 0.472 |
| MIRNA_MM_GRIMSON_MIR-486-5P | 121 | 0.426 | 1.457 | 0 | 0.308 | 0.701 |
| MIRTARBASE_MM_HSA-MIR-29C | 20 | 0.894 | 1.449 | 0.034 | 0.312 | 0.737 |

**Fig. 2.**
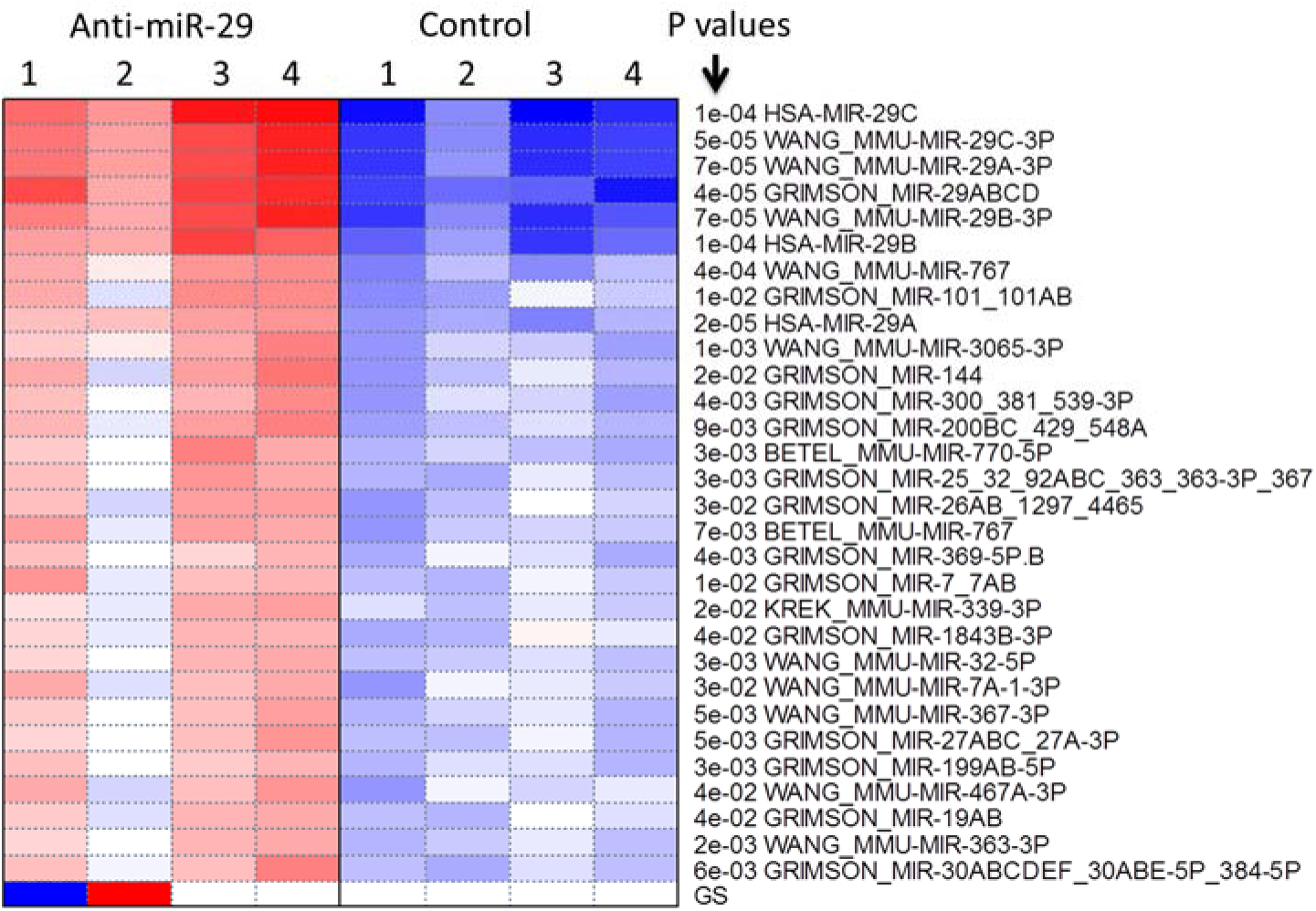
Significant gene sets from PGSEA analysis. Red indicates higher expression of genes targeted by certain microRNA according to prediction, while blue means lower expression.

To gain further insight into the downstream molecular pathways, we analyze this expression data using other gene sets in GSKB. Using the PGSEA software, we identified significantly altered pathways in many different types of gene sets. Table 4 shows some of the top ranked gene sets by category. In the co-expression category, we detected that the change in expression profile when miR-29c was suppressed are similar to several reported expression signatures. The top 5 signatures are related to various perturbations of liver cells such as overexpression of TCFAP2C [18], or treatment with rosiglitazone [19], a compound that can lower glucose levels. The third most significantly gene set (Nones_Mdr1A_Curcumin-Diet_Fibrogenesis_Dn) includes genes involved in fibrogenesis down-regulated in the colon of multidrug resistance gene-deficient (mdr1a-/-) mice fed with curcumin diet [20]. Fibrogenesis is a process regulating the deposition of extracellular matrix proteins. This is in fact a reoccurring theme in multiple significant gene sets across categories. For example, two significant Gene Ontology terms are “Extracellular Matrix Structural Constituent”, and “Collagen Fibril Organization”. Other related gene sets are “ECM-Receptor Interaction”, “Integrin”, and “Beta Integrin Cell Surface Interactions”. These results strongly suggest that lower expression of miR-29 caused by antisense oligonucleotides leads to the up-regulation of extracellular matrix proteins and fibrogenesis in hepatic cells. This agrees with the well-established role of miR-29 in fibrosis in liver [21], and several other tissues/cells such as heart [22], stellate cells [23] and HK-2 cells (human kidney cell line) [24] *etc*. All members of the miR-29 family were found to be downregulated in murine livers treated with carbon tetrachloride to induce hepatic fibrogenesis [21]. Roderburg et al. [21] also noted abnormally low expression of miR-29 in patients with advanced liver fibrosis and liver cirrhosis. Thus GSKB enables us to identify downstream molecular pathways regulated by the miR-29 family.

**Table 4.** Top gene sets in all categories from PGSEA analysis of the anti-miR-29 treated liver cells.

| Top Gene Sets by category | Pval | Diff. in T statistics |
| --- | --- | --- |
| **Co-Expression** |  |  |
| Wang_Rosiglitazone_Adipocyte-Secreted-Protein_Diff | 8.30E-04 | 14.9 |
| Holl_Tcfap2C_Liver_Dn | 2.12E-04 | -15.6 |
| Nones_Mdr1A_Curcumin-Diet_Fibrogeneis_Dn | 2.44E-04 | 15.4 |
| Burke_Transcriptional-Profile_Liver-Only_Dn | 2.78E-04 | -15.2 |
| Rampon_Pcdh12_Placenta_Cell-Matrix-Adhesion_Migration_Diff | 4.83E-04 | 13.6 |
| **Gene Ontology** |  |  |
| Extracellular_Matrix_Structural_Constituent | 2.22E-04 | 18.9 |
| Collagen_Fibril_Organization | 7.30E-04 | 12.9 |
| Cellular_Response_To_Amino_Acid_Stimulus | 4.62E-04 | 12.1 |
| Glutathione_Transferase_Activity | 5.26E-04 | -11.1 |
| Basement_Membrane | 9.15E-04 | 9.3 |
| **Metabolic Pathways** |  |  |
| Cd44_Pathway | 3.11E-04 | 16.9 |
| Ecm-Receptor_Interaction | 4.00E-04 | 11.4 |
| Protein_Digestion_And_Absorption | 1.11E-04 | 10.9 |
| Cav1_Pathway | 5.73E-04 | 9.1 |
| Glutathione_Metabolism | 4.51E-05 | -9.2 |
| **Mirna Target Genes** |  |  |
| Hsa-Mir-29C | 1.45E-04 | 17.3 |
| Wang_Mmu-Mir-29C-3P | 5.19E-05 | 15.2 |
| Wang_Mmu-Mir-29A-3P | 7.38E-05 | 15.0 |
| Grimson_Mir-29Abcd | 3.91E-05 | 15.6 |
| Wang_Mmu-Mir-29B-3P | 7.32E-05 | 14.8 |
| **Currated Pathways** |  |  |
| Integrin | 2.59E-04 | 8.4 |
| Pyruvate_Metabolism | 9.66E-04 | -4.9 |
| **Transcription Factor Target Genes** |  |  |
| Srebf1 | 4.62E-04 | -7.9 |
| Smad3 | 4.01E-04 | 7.1 |
| **Other gene sets** |  |  |
| Beta1_Integrin_Cell_Surface_Interactions | 2.77E-04 | 13.7 |
| Mitral_Valve_Prolapse | 9.29E-04 | 11.8 |
| Abnormality_Of_The_Mitral_Valve | 8.73E-04 | 11.8 |
| Syndecan-1-Mediated_Signaling_Events | 5.21E-04 | 11.6 |
| Osteoarthritis | 8.89E-04 | 11.0 |

Table 4 also shows transcription factors that might be involved. Treatment with anti-miR-29 oligonucleotides is associated with reduced expression of *Srebf1* (sterol regulatory element binding transcription factor 1) target genes. Srebf1 is regulator of lipid homeostatsis [25] and directly binds to sterol regulatory element-1 (SRE1) flanking LDL receptor genes and other related genes. This suggests that lower expression of miR-29 might hinder the normal function of hepatic cells in metabolism. Therefore high expression of miR-29 expression in hepatic cells is required for lipid metabolism and that Srebf1 might be downstream of miR-29. There does not appear to be any existing evidence linking miR-29 with Srebf1. This is a directly testable hypothesis for further experimental studies regarding the function of miR-29.

Table 4 also indicates that anti-miR-29 treatment lead to increased expression of *Smad3* (SMAD family member 3) target genes. It is well-established that Smad3 is a key player in TGFβ-mediated fibrosis, tumor suppression and metastasis [26]. Qin et al. provided evidence that Smad3-mediated suppression of miR-29 expression by *TGFβ1* is achieved by direct binding to the promoter of miR-29 [27], and overexpression of miR-29b inhibits the collagen I and III and prevents renal fibrosis. In our analysis, suppression of miR-29 leads to the upregulation of Smad3 target genes, suggesting that Smad3 and miR-29 might form a negative regulation loop. Indeed, Xiao et al. [28] provided some evidence that gene transfer of miR-29 was able to block bleomycin-induced pulmonary fibrosis by suppressing the expression of TGFβ-1and inhibiting Smad3 phosphorylation.

To further confirm these findings, we analyzed another expression data set (GSE27035), in which fetal astrocytes were transfected with miR-29 [29]. We obtained similar results (data not shown) regarding the extracellular matrix related gene sets. For transcriptional factors, E2F family members are highly significant, which is not observed in hepatic cells in the previous dataset. But Smad3 targets genes are downregulated, suggesting that the universal regulation of extracellular matrix genes by miR-29 might be related to Smad3.

### Identifying transcription factors from gene expression data

In another example, we used our knowledgebase to detect transcription factors (TFs) responsible for tissue-specific gene expression. We analyzed a large gene expression dataset consisting of 3 or 4 biological replicates for each of the 24 mouse tissues [30] (NCBI accession number GSE24207). We used a subset of 373 predicted TF target gene sets in our analysis using PGSEA. Fig. 2 lists top 30 most significant TFs associated with various tissues. Many highly significant TFs are known to be involved in different organs. The most significant are the hepatocyte nuclear factors (HNF4A, HNF1A, and FOXA2/HNF3B) that are highly expressed in the liver and are supported by many studies to be involved in liver development [31]. The target genes of SPI1 (Spleen focus forming virus proviral integration oncogene, also listed as SFPI1) are highly expressed in spleen and bone marrow, in agreement with the fact that SPI1 is an ETS-domain transcription factor involved in myeloid and B-lymphoid cell development [32]. Another very highly significant TF, Steroidogenic Factor-1 (NR5A1), is found to be responsible for adrenal gland-specific expression profile. This nuclear receptor is known to play an important role in adrenal development and function and mutations in this protein are associated with adrenal hypolasia [33]. PPARG (peroxisome proliferator activated receptor gamma) is essential for the differentiation of adipose tissue [34]. In our result, its target genes were found to be significant highly expressed in the adipose tissue. Most of the TFs in Fig. 2 are supported by previous studies in the literature.

### Meta-analysis of published gene lists yields insight into blindness and visual perception

The comprehensive dataset can also serve as an information source on published expression signatures. For example, using a keyword “retina” to conduct a search at our web site, we retrieved 56 published signatures from 26 genome-wide expression studies of the retina cells. Detailed information on each of gene lists, including links to PubMed, is available for further examination.

Meta-analysis of a set of retrieved signatures can provide insights into the relationship among multiple previously published expression profiles. We conducted an all-versus-all overlapping analysis of these 56 gene lists. Following the method developed previously [35], we generated a network where nodes correspond to gene lists and edges represent significant overlaps between them (Fig.4). We found 52 significant overlaps (FDR < 0.0001) among 25 gene lists. Interestingly, the overlaps define two groups with high similarity within each group and very little in between. Group A on the left side of Fig. 4 includes gene lists that are upregulated in ageing, bright-light-damaged retina, or other injury, as well as hypoxia treatment from 3 independent studies[36–38]. Retinal hypoxia is believed to be the mechanism of blinding underlying several diseases [39] and have been subjected to studies using animal models. Our results suggest that there is significant similarity in gene expression response across multiple studies. We identified the most frequently shared genes across these 10 gene lists in group A. Table 5 lists 29 genes that are shared by 3 or more gene lists in group A gene lists. These 29 genes are significantly enriched in genes related to inflammatory response (P value < 1.80E-05), and immune response (P value < 5.30E-03). This is in agreement with previous finding regarding the activation of inflammatory response upon hypoxia treatment [39]. As shown in Table 5, the most commonly upregulated genes in group A is the glial fibrillary acidic protein (GFAP), which was initially discovered as an indicator of stress in astrocytes in the brain, but its activation in the radial glia (Müller cells) is also known to signal stress in the retina [40]. Based on the consensus of 8 genome-wide expression studies, our new gene lists in Table 5 thus could serve as marker for stress response in the retina caused by hypoxia or other injuries.

**Fig. 3.**
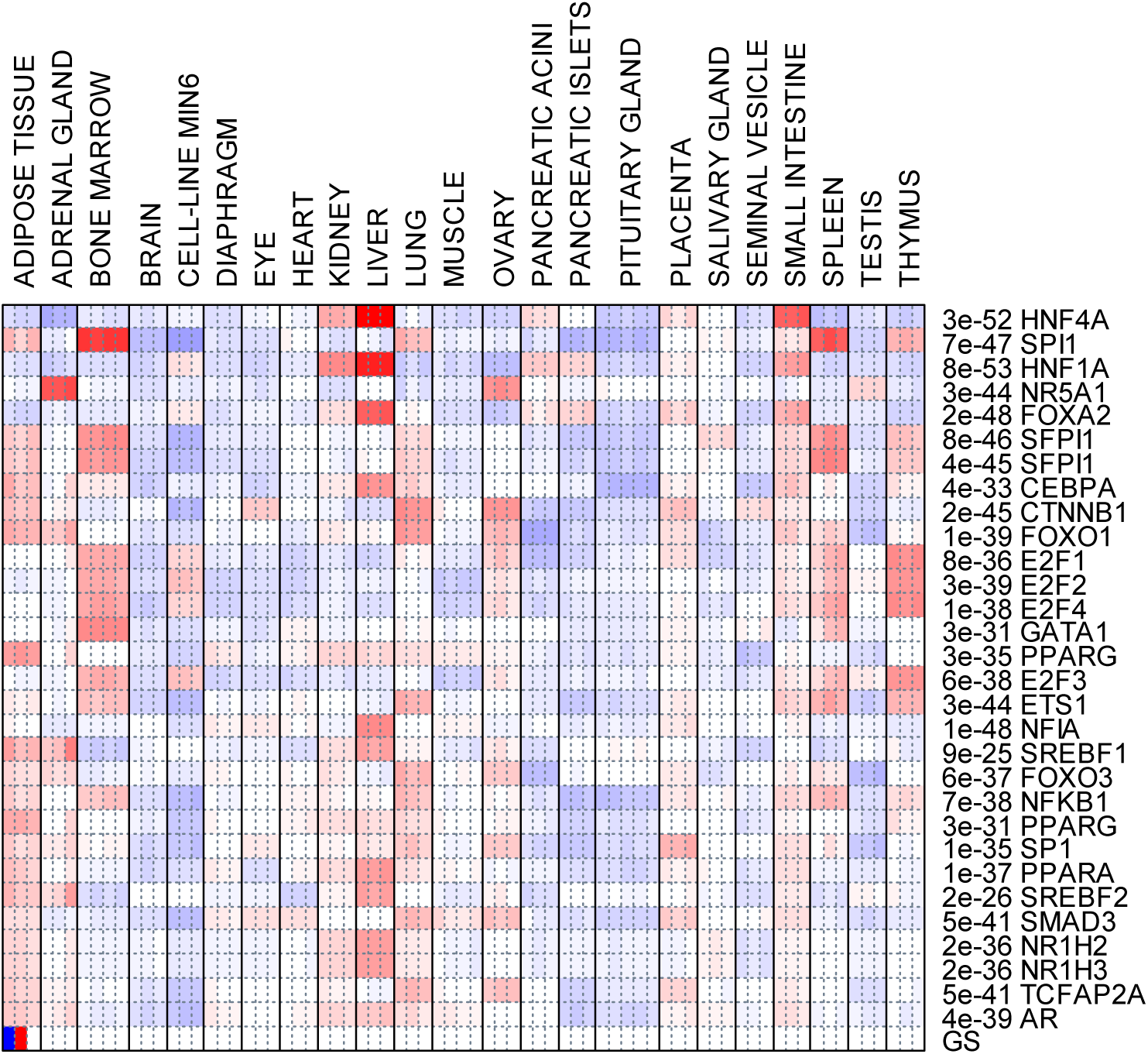
Significantly altered gene sets in normal tissues. Red represents higher expression of a set of genes regulated by a transcription factor.

**Fig. 4.**
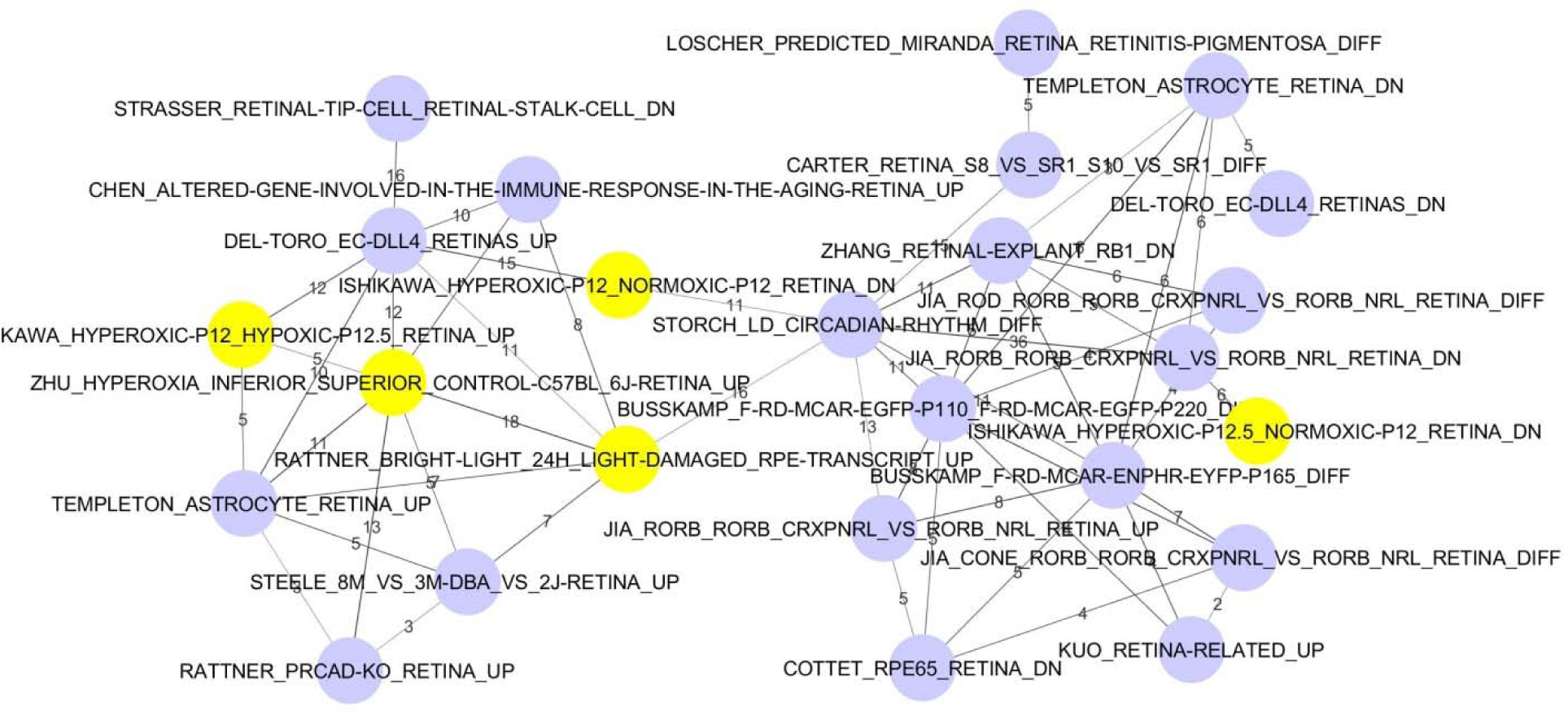
Overlapping analysis of 46 published lists of differentially expressed genes related to the retina. Each note represents a gene list. Edges represent significant overlaps. Edge labels are the number of genes shared and the width indicates statistical significance. Highlighted in yellow are lists related to hypoxia.

**Table 5.**
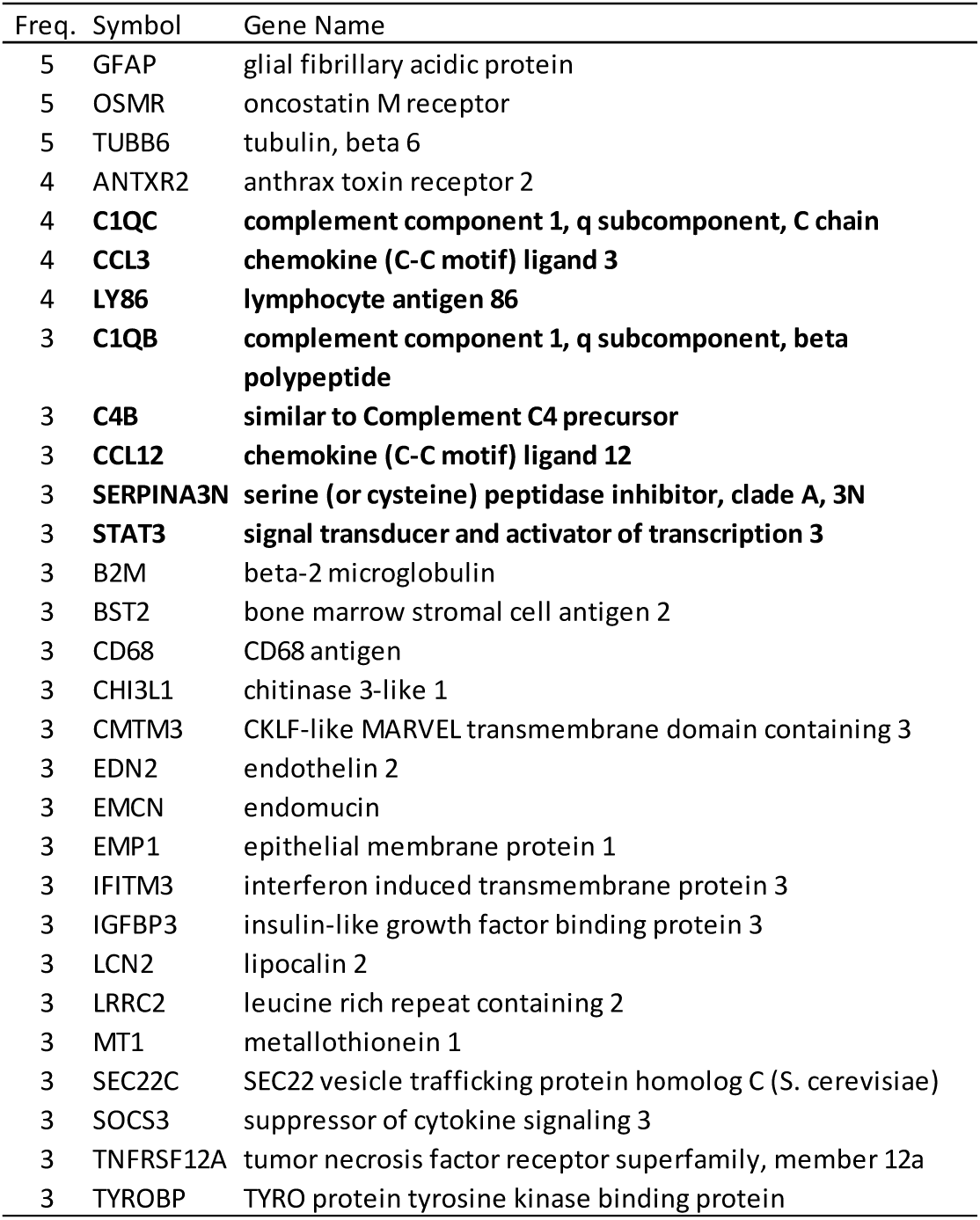
Genes frequently appeared in gene set cluster A. Genes in bold are related to inflammatory response. Freq.: frequency, the number of gene sets with the gene.

On the other hand, the 15 gene lists in group B on the right side of Fig. 4 have different biological theme. Three of the gene lists (ZHANG_RETINAL-EXPLANT_RB1_DN, COTTET_RPE65_RETINA_DN, DEL-TORO_EC-DLL4_RETINAS_DN) are genes downregulated in retina with mutated genes (Rb1[41], RPE65[42], and DLL4[43]), compared with normal retina. Another gene list includes genes downregualted in hypoxic retina [36]. This seems to suggest that gene lists in this group might include genes specifically required for normal photoreceptive function of the retina cells. This is confirmed by examining the frequently appearing genes in this group. Among the 38 genes (Table 6) that are shared by 3 or more lists in this group, half of them are related to visual perception according to GO, which is extremely significant (P < 1.2E-28). The most frequently appearing gene ARR3 (arrestin 3, retinal) are predicted to play an important role in retina-specific signal transduction with possible binding to photoactivated-phosphorylated opsins, including OPN1SW (opsin 1) that are shared by 7 of the 15 gene lists (Table 6). Based on multiple studies, table 6 serves as a reliable list of retina-specific genes important for photoreception.

**Table 6.** Genes frequently appeared in gene sets cluster B. Genes in bold are related to visual perception according to GO.

| Freq. | Symbol | Gene Name |
| --- | --- | --- |
| 8 | **ARR3** | **arrestin 3, retinal** |
| 7 | **OPN1SW** | **opsin 1 (cone pigments), short-wave-sensitive (color blindness, tritan)** |
| 6 | **CNGA1** | **cyclic nucleotide gated channel alpha 1** |
| 6 | **GNAT2** | **guanine nucleotide binding protein, alpha transducing 2** |
| 6 | **PDE6B** | **phosphodiesterase 6B, cGMP, rod receptor, beta polypeptide** |
| 6 | **PDE6H** | **phosphodiesterase 6H, cGMP-specific, cone, gamma** |
| 5 | **GNAT1** | **guanine nucleotide binding protein, alpha transducing 1** |
| 5 | **PDE6C** | **phosphodiesterase 6C, cGMP specific, cone, alpha prime** |
| 5 | **RCVRN** | **recoverin** |
| 5 | GNB3 | guanine nucleotide binding protein (G protein), beta 3 |
| 5 | EA2 | erythrocyte antigen 2 |
| 4 | **PDE6G** | **phosphodiesterase 6G, cGMP-specific, rod, gamma** |
| 4 | **RPGRIP1** | **retinitis pigmentosa GTPase regulator interacting protein 1** |
| 4 | AQP1 | aquaporin 1 |
| 4 | CRX | cone-rod homeobox containing gene |
| 4 | GNGT2 | guanine nucleotide binding protein (G protein), gamma transducing activity polypeptide 2 |
| 4 | NR2E3 | nuclear receptor subfamily 2, group E, member 3 |
| 3 | **ABCA4** | **ATP-binding cassette, sub-family A (ABC1), member 4** |
| 3 | **CNGA3** | **cyclic nucleotide gated channel alpha 3** |
| 3 | **CNGB3** | **cyclic nucleotide gated channel beta 3** |
| 3 | **GUCA1B** | **guanylate cyclase activator 1B** |
| 3 | **OPN1MW** | **opsin 1 (cone pigments), medium-wave-sensitive (color blindness, deutan)** |
| 3 | **PDE6A** | **phosphodiesterase 6A, cGMP-specific, rod, alpha** |
| 3 | **ROM1** | **rod outer segment membrane protein 1** |
| 3 | **SLC24A1** | **solute carrier family 24 (sodium/potassium/calcium exchanger), member 1** |
| 3 | CALB2 | calbindin 2 |
| 3 | CRYBA1 | crystallin, beta A1 |
| 3 | GAS7 | growth arrest specific 7 |
| 3 | GLB1L3 | galactosidase, beta 1 like 3 |
| 3 | GNGT1 | guanine nucleotide binding protein (G protein), gamma transducing activity polypeptide 1 |
| 3 | MEF2C | myocyte enhancer factor 2C |
| 3 | NEFM | neurofilament, medium polypeptide |
| 3 | NRL | neural retina leucine zipper gene |
| 3 | NXNL2 | nucleoredoxin-like 2 |
| 3 | PDGFRA | platelet derived growth factor receptor, alpha polypeptide |
| 3 | PITPNM3 | PITPNM family member 3 |
| 3 | SPC25 | SPC25, NDC80 kinetochore complex component, homolog (*S. cerevisiae*) |
| 3 | TRPM1 | transient receptor potential cation channel, subfamily M, member 1 |

Overall, through meta-analysis of 56 retina-related gene lists, we revealed two fundamentally different types of gene expression changes in the retina. One is related to stress response and inflammatory response that are blamed for blindness in many diseases; the other is a set of genes that are required by visual perception by the retina cells. Our database could be used to conduct similar meta-analysis using various search keywords.

## CONCLUSION

We have created a comprehensive gene set database for pathway analysis in mouse. We also demonstrated that this database could be used with different pathway analysis software to gain insights into genome-wide expression profiles. For further improvement of knowledgebase, we will update the database from existing sources, and also continue to improve the accuracy of the existing curation, and search for additional published gene lists.

## ACKNOWLEDGEMENTS

The authors thank Drs. Jill Mesirov and Arthur Liberzon for advices on gene list collection, and the reviewers for many constructive suggestions

